# High-Throughput Screening on Primary Tumor-Associated Microglia and Macrophages Identifies HDAC Inhibitors as Enhancers of Phagocytosis and Potent Partners for Immunotherapy in Glioblastoma

**DOI:** 10.1101/2025.03.24.645104

**Authors:** Mona Khalaj, Madison L. Gutierrez, Parisa Nejad, Tal Raveh, Faranak Fattahi, Irving L. Weissman

## Abstract

Glioblastoma multiforme (GBM) is a lethal brain tumor with limited treatment options. Tumor-associated macrophages and microglia (TAMs) drive immune suppression and tumor progression, making them a key therapeutic target for GBM. Enhancing TAM phagocytosis in GBM has shown promise, particularly with innate checkpoint inhibitors, such as CD47-blocking antibodies. However, small molecule approaches, which offer tunable and potentially synergistic mechanisms, remain underexplored in this context. In this study, we conducted the first large-scale chemical screen on primary TAMs from patients with GBM, identifying histone deacetylase (HDAC) inhibitors as potent inducers of phagocytosis. These compounds demonstrated phagocytosis-inducing effects across multiple GBM patient samples, with further amplification when combined with CD47 blockade. In a xenograft GBM model, HDAC inhibitors enhanced phagocytosis and suppressed tumor growth, with even greater efficacy in combination with CD47 antibodies. Our findings highlight HDAC inhibitors as promising agents to reprogram TAMs and synergize with immune checkpoint therapies, offering a novel strategy to bolster anti-tumor immunity in GBM.

## Introduction

Glioblastoma multiforme (GBM) is an aggressive and universally lethal primary brain tumor with limited treatment options and a median survival of approximately 15 months despite standard-of-care treatments, including surgery, radiation, and temozolomide chemotherapy (1). The tumor microenvironment (TME) of GBM is dominated by tumor-associated macrophages and microglia (TAMs), which often adopt an immunosuppressive phenotype that supports tumor growth and resistance to therapy. Therefore, reprogramming TAMs to enhance their anti-tumor function represents an exciting therapeutic strategy (2,3).

In the healthy brain, central nervous system–resident immune cells primarily consist of microglia, which are yolk sac–derived myeloid cells (4). In the context of brain tumors, these resident cells are joined by infiltrating monocytes that differentiate into macrophages. We have previously shown in a mouse model that both resident microglia and infiltrating macrophages act as effector cells capable of phagocytic clearance of GBM cells upon CD47 blockade (5). In this study, we collectively refer to these cells as tumor-associated microglia and macrophages (TAMs).

Phagocytosis is a critical function of macrophages that clears cancer cells and activates anti-tumor adaptive immune responses. Strategies to enhance macrophage function have led to the development of innate immune checkpoint inhibitors, such as blocking antibodies against CD47, LILRB1, PD-1 and CD24, which override the ‘don’t eat me’ signal and promote cancer cell engulfment (6–11). Although these therapies have shown promise in preclinical studies and early clinical trials, their efficacy varies among patients, highlighting the need for additional approaches to enhance cancer cell clearance via TAM phagocytosis.

Genome-wide functional CRISPR screens have uncovered key genes involved in macrophage phagocytosis and inflammatory pathways (12–14), primarily using myeloid cell lines (e.g., U937 or J774 cells) or PBMC-derived macrophages . While these models have provided critical mechanistic insights, they do not recapitulate the complexity of the tumor microenvironment (TME) in which TAMs operate. Given the challenges of performing large-scale genetic screens in primary TAMs, small molecule-based approaches offer a practical strategy. In this work, we present the first high-throughput functional screen conducted on TAMs isolated from primary patient samples, enabling the identification of small molecules that modulate phagocytosis in a clinically relevant manner.

To discover small molecules that potently activate TAM phagocytosis, we conducted a screen using a library of 1,377 FDA-approved pharmaceutical compounds on primary patient-derived TAMs. Among the top hits, histone deacetylase (HDAC) inhibitors, including Pracinostat and Resminostat, emerged as potent inducers of phagocytosis, demonstrating dose-dependent activity in patient-derived TAM samples. These findings align with prior reports of HDAC inhibitors modulating macrophage function in infectious diseases and inflammation (15–19).

We have previously demonstrated that CD47 antibodies effectively promote GBM tumor clearance by inducing phagocytosis (5,20,21). Notably, this study further revealed a synergistic effect between HDAC inhibitors and CD47-blocking antibodies, highlighting a combination strategy to overcome immune suppression in GBM. Preclinical validation in a xenograft GBM model demonstrated robust anti-tumor activity with HDAC inhibition alone, which was further enhanced in combination with CD47 blockade. This work underscores the translational potential of integrating HDAC inhibitors with immunotherapies (22). By systematically investigating the role of HDAC inhibitors in TAM reprogramming and phagocytosis, this study contributes to the development of novel therapeutic strategies aimed at addressing the profound immune suppression characteristic of GBM.

## Results

### High-Throughput Chemical Screen Identifies Phagocytosis-Inducing Drugs

To identify pharmaceutical compounds capable of enhancing macrophage phagocytosis in GBM, we performed a high-throughput chemical screen on human TAMs freshly isolated from resected primary GBM tumors. CD11b+ microglia and macrophages were enriched from patient surgical resections and treated overnight with a library of 1365 FDA-approved compounds. Following treatment, GFP-labeled patient-derived GBM cells were used in a phagocytosis assay to quantify phagocytic activity (**Figure 1A, Figure S1**). Compounds that induced phagocytosis at levels greater than 2 standard deviations above the mean were classified as significant hits. A scatter plot of the Z-scores for all tested compounds illustrates the distribution of activity, with significant hits clearly separated from the majority of inactive compounds (**Figure 1B**).

**Figure 1.**
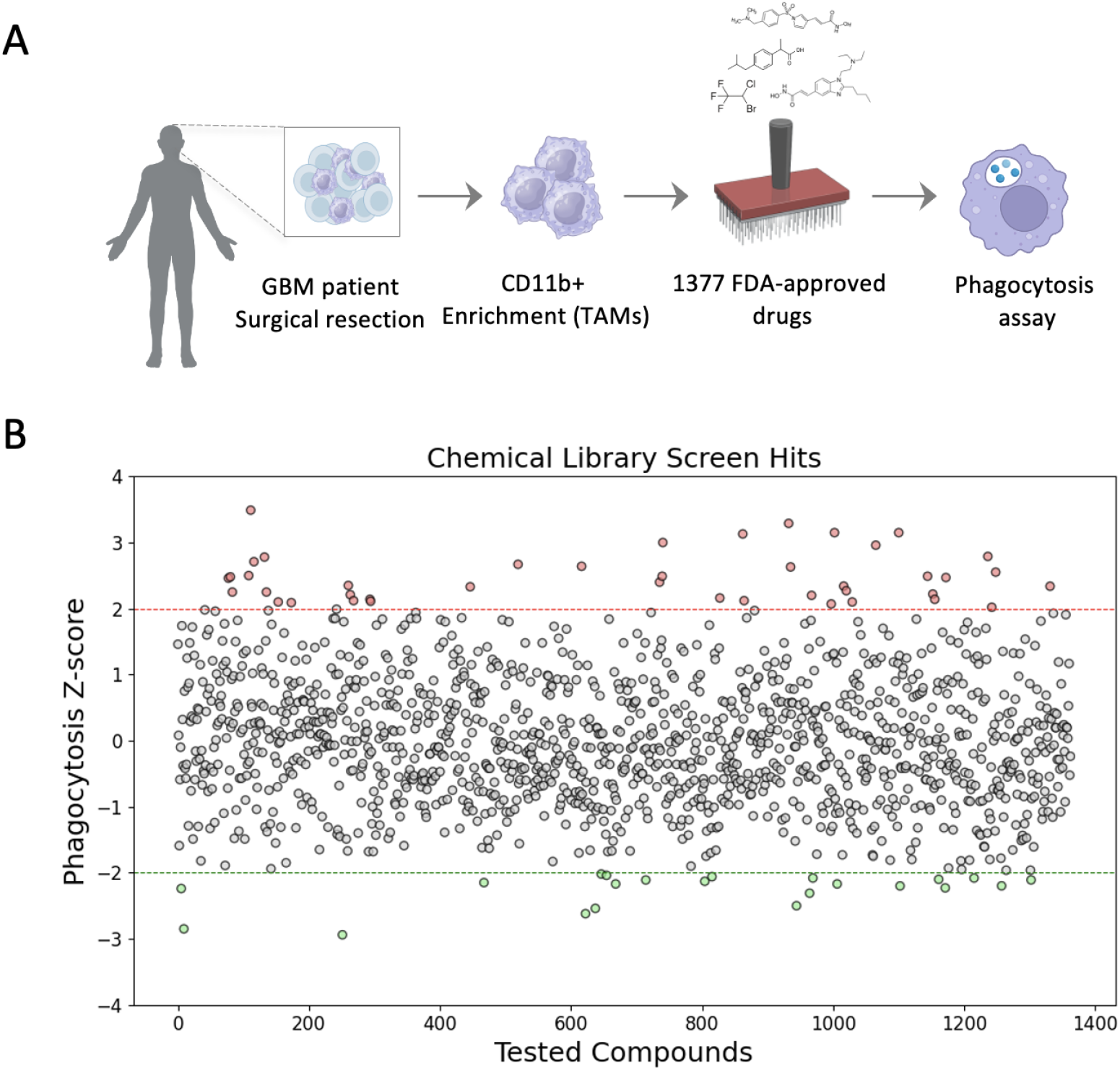
High-Throughput Chemical Screen on TAMs from GBM Patients Identifies Drugs that Induce Phagocytosis. **A**. Schematic of the chemical screen workflow: 1365 FDA-approved drugs were tested on tumor-associated macrophages (TAMs) isolated from GBM patient surgical resections. CD11b+ TAMs were enriched, treated and subjected to a phagocytosis assay to identify compounds that enhance phagocytic activity. **B**. Scatter plot of the Z-scores for all tested compounds. Z-scores represent the number of standard deviations from the mean, with compounds inducing phagocytosis at >2 standard deviations above the mean considered significant hits.

### HDACs Are Enriched Among Phagocytosis-Inducing Targets

To identify top targets from the screen, we applied multiple statistical approaches to assess target enrichment among compounds that significantly impact phagocytosis. We obtained isomeric SMILES (Simplified Molecular Input Line Entry System) representations for each drug in the FDA-approved library from Selleckchem and used the Similarity Ensemble Approach (SEA) computational tool (23) to predict drug-protein interactions. Predicted interactions were filtered to retain only human proteins with a significance threshold of p < 0.05.

To quantify target enrichment, we calculated weighted combined z-scores by summing normalized z-scores across all compounds targeting a given protein (23). This analysis identified HDACs as one of the most enriched target classes among compounds with significant effects on phagocytosis (**Figure 2A**).

**Figure 2.**
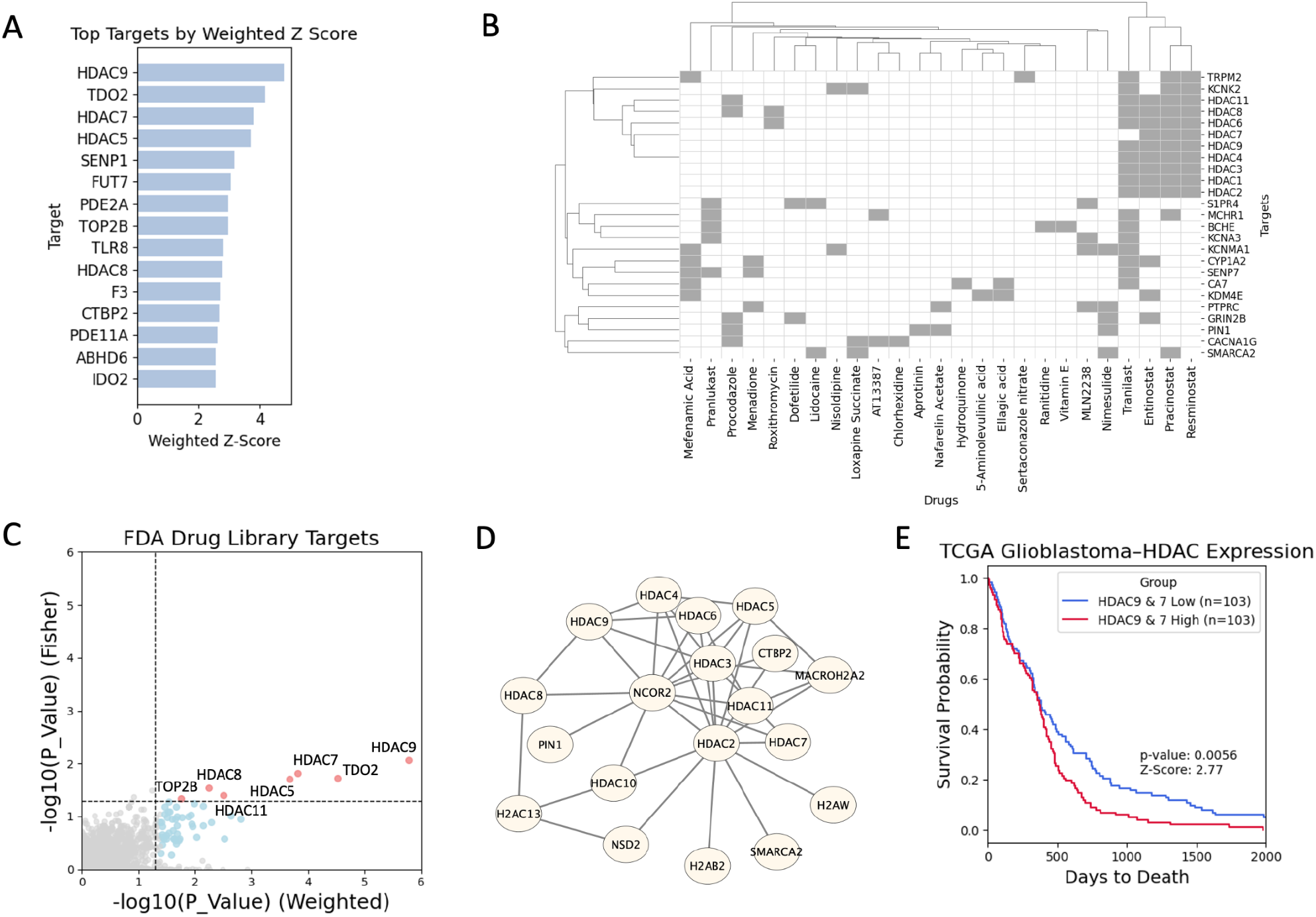
HDACs Are Significantly Enriched Among Targets Identified in the Chemical Screen. **A**. Enriched target genes were identified along with their combined Z-scores, calculated by integrating the normalized Z-scores from all treatments associated with each gene. This combined Z-score approach provides a comprehensive measure of the significance of gene enrichment across multiple treatments. **B**. A drug-gene interaction matrix was generated for the 25 most frequently-enriched target genes within hit drugs with Z score >2 identified in Figure 2A. This matrix highlights the interactions between drugs and their associated target genes, offering insights into key drug-target relationships. **C**. As an independent validation, a Fisher’s exact test was performed to assess enrichment of targets in drugs with Z-scores > 2, while also identifying depletion of these targets in drugs with Z-scores < 2. This secondary analysis strengthens confidence in the robustness of the enriched target genes. **D**. To further analyze these findings, a STRING protein-protein interaction (PPI) network was constructed, mapping interactions between the top hits from the combined Z-score analysis. This network visualization provides a deeper understanding of functional connections between enriched targets. **E**. Kaplan-Meier survival analysis of glioblastoma patients stratified by the expression of top two hit targets, HDAC9,7. Patients were divided into two groups based on HDAC9,7 expression levels: High expressors (top 25%; HDAC9,7 High) shown in red and Low expressors (bottom 25%; HDAC9,7 Low) shown in blue. The analysis demonstrates a significant difference in survival probability between the two groups, with low HDAC9,7 expression associated with improved survival (log-rank test, p = 0.0056).

Building on these findings, we constructed a drug-gene interaction matrix by integrating SEA-derived drug-target data to map key relationships between enriched targets and their associated compounds (**Figure 2B**). This analysis revealed a dominant cluster of HDAC-targeting compounds, highlighting their potential role in modulating TAM phagocytosis. Among these, Resminostat and Pracinostat were prioritized for further validation based on their selective HDAC targeting and minimal off-target interactions.

To further validate the significance of HDACs as key regulators of TAM activity, we performed a Fisher’s exact test to assess the enrichment of HDACs among compounds with enrichment scores >2 and their depletion among compounds with enrichment scores <2. This analysis confirmed that HDAC-targeting compounds were significantly overrepresented among the hits, underscoring their centrality in the regulation of TAM phagocytosis (**Figure 2C**). To further dissect functional relationships, we generated a STRING protein-protein interaction (PPI) network, which positioned HDACs as central nodes among top-enriched targets (**Figure 2D**), further emphasizing their pivotal role in this context. Together, these analyses robustly establish HDACs as major therapeutic targets for modulating TAM function in GBM.

### Functional validation of HDAC Inhibitors in GBM Patient-Derived TAMs

As mentioned above, we prioritized Pracinostat and Resminostat for further evaluation due to their strong phagocytosis-enhancing activity and specificity for HDAC targets. To assess their effects, TAMs were isolated from a second GBM patient and treated with these compounds at increasing doses. Both Pracinostat and Resminostat demonstrated dose-dependent increases in phagocytic activity, as quantified by GFP-labeled tumor cell uptake, with significant effects observed at concentrations of 15 µM and 45 µM (**Figures 3A-B**).

**Figure 3.**
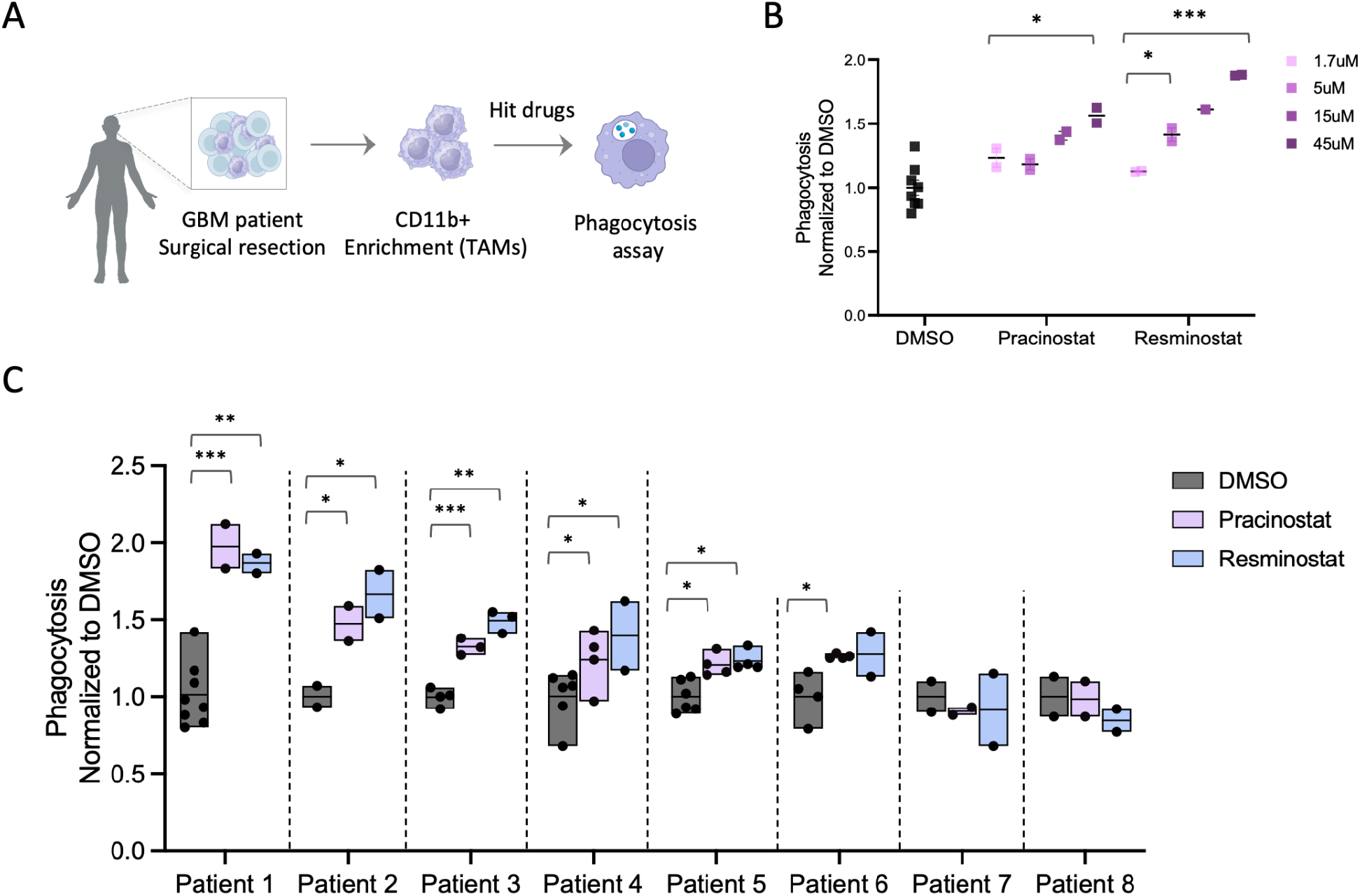
Validation of HDAC inhibitors Resminostat and Pracinostat for Stimulating Phagocytosis by TAMs from Multiple GBM Patient Samples. **A**. Schematic of the experimental workflow. TAMs were enriched from GBM patient surgical resections using CD11b-positive selection. Enriched TAMs were treated overnight hit compounds (Resminostat and Pracinostat) or DMSO (control). The following day, TAMs were co-incubated for 2 hours with GFP-tagged primary GBM tumor cells, and phagocytosis was assessed. **B**. Dose-dependent effects of Resminostat and Pracinostat on phagocytosis in TAMs from a second GBM patient sample. TAMs were treated with increasing concentrations of Resminostat and Pracinostat (1.7 µM, 5 µM, 15 µM, and 45 µM) and then incubated with GFP-tagged primary GBM cells. Phagocytosis was quantified and normalized to the DMSO control. Both compounds demonstrated a dose-dependent increase in phagocytosis. **C**. Phagocytosis activity of TAMs across eight GBM patient samples. TAMs from individual patients were treated with 5µM to 15µM Resminostat, 5µM to 15µM Pracinostat, or DMSO control overnight, followed by a 2-hour incubation with GFP-tagged GBM cells. Phagocytosis was quantified and normalized to the DMSO control for each patient. Data are shown as mean ± SEM.

To confirm the generalizability of these findings, we tested both compounds across TAMs isolated from eight independent GBM patient samples. Pracinostat and Resminostat consistently enhanced phagocytosis in a majority of the tested samples (six out of eight patients), underscoring their potential as broadly applicable therapeutic agents for GBM (**Figure 3C**).

### Synergy Between HDAC Inhibitors and CD47 Blockade

Given the growing interest in immune checkpoint inhibitors, we investigated whether HDAC inhibitors could synergize with CD47 blockade. TAMs from three GBM patients were treated with Pracinostat or Resminostat in combination with a CD47-blocking antibody (5F9) (**Figure 4A**). Phagocytosis assays revealed that both compounds significantly enhanced the effects of CD47 blockade, with a marked increase in GFP-labeled tumor cell uptake compared to either agent alone (**Figure 4B,C** & **Figure S2**). These results suggest that combining HDAC inhibitors with CD47-targeted therapies could amplify therapeutic efficacy in GBM.

**Figure 4.**
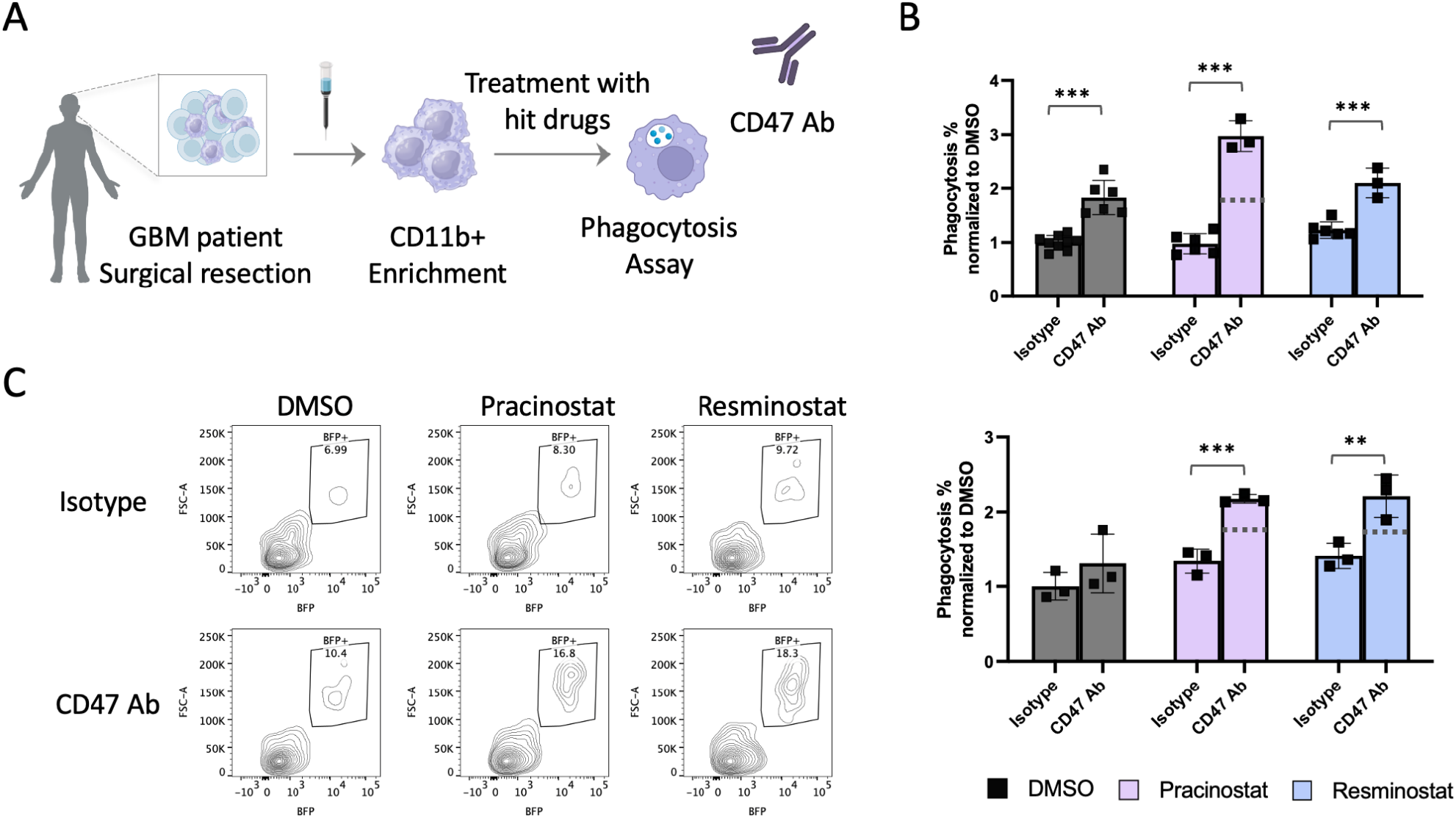
Synergistic Effects of Pracinostat and Resminostat with CD47 Antibody in Enhancing Phagocytosis. **A**. Schematic of the experimental setup to test the synergy of hit compounds Pracinostat and Resminostat with CD47 blockade. TAMs isolated from primary GBM samples were treated overnight with the drugs, followed by a 2-hour phagocytosis assay in the presence of either CD47-blocking antibody (5F9) or an isotype control. **B**. Phagocytosis results from Three GBM patients normalized to the DMSO control. Both Pracinostat and Resminostat significantly enhance phagocytosis in the presence of CD47 antibody compared to the isotype control, demonstrating a synergistic effect. Data points represent individual technical replicates, and horizontal bars denote the mean. Dashed lines indicate phagocytosis levels assuming additive effects. **C**. Representative flow cytometry plots of a primary TAM sample showing synergy between Pracinostat and CD47 Abs. Data are shown as mean ± SEM.

### Validation in a Mouse Model of GBM

To assess the translatability of our findings, we utilized a mouse model of GBM by stereotactic injection of CT2A mouse GBM cells into the brains of C57BL/6 mice (**Figure 5A**). TAMs were isolated from murine tumors and treated with Pracinostat and Resminostat *in vitro*. Both compounds induced dose-dependent increases in phagocytosis, mirroring the effects observed in patient-derived TAMs (**Figure 5B**).

**Figure 5.**
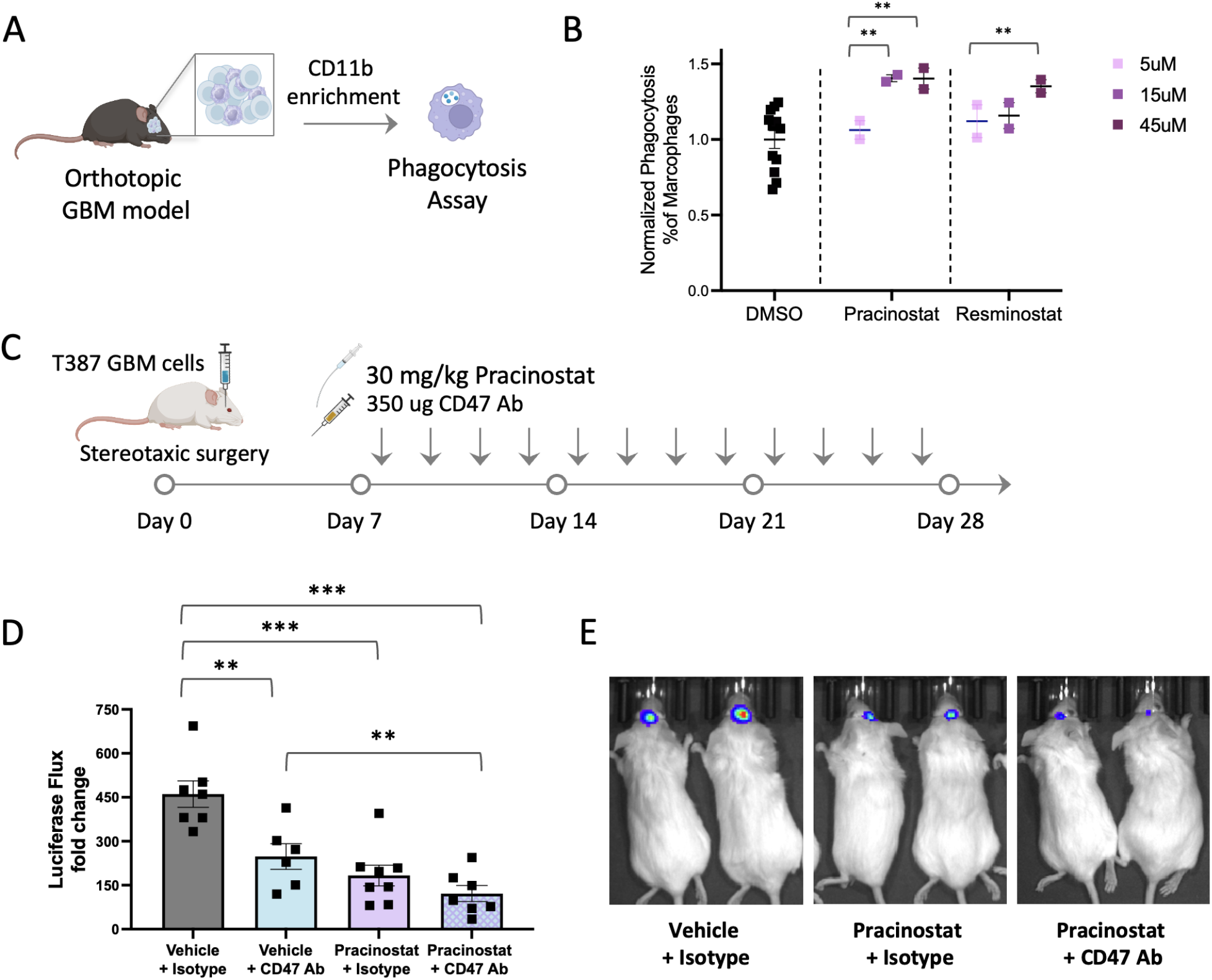
Validation of Hit Compounds in a Mouse Model of GBM. **A**. Schematic of the experimental workflow: TAMs were isolated from stereotactic CT2A GBM mouse models via CD11b+ enrichment. The phagocytic activity of TAMs was assessed after treatment with HDAC inhibitors (Pracinostat, Resminostat). **B**. Dose titration results showing normalized phagocytosis in TAMs isolated from the mouse model. **C**. Schematic for the treatment of GBM xenografts to assess the impact of hit compounds on tumor growth with or without CD47 antibody blockade. 100K BFP+ Luciferase+ T387 human GBM cells were injected stereotactically into the brain, and mice were treated with oral Pracinostat (30 mg/kg) and subcutaneous CD47 Ab (350 µg) four times a week. **D**. Quantification of tumor burden based on normalized luciferase signal at Day 22, normalized to the start of treatment (day 7). **E**. Representative bioluminescence imaging of tumor burden at Day 15 post-injection, comparing Vehicle + Isotype (left) with Pracinostat + Isotype (middle) and Pracinostat + CD47 Ab combination therapy (right), indicating decreased tumor burden. Data are shown as mean ± SEM.

To further evaluate the in vivo efficacy of these compounds, we assessed their impact on tumor progression in xenograft models of GBM. Human T387 cells were stereotactically injected into the brains of NSG mice to establish the tumors. Given that Pracinostat has a higher predicted blood-brain barrier permeability (Pracinostat LogP: 2.57 vs. Resminostat LogP: 1.1), we prioritized it for this experiment. Analysis of fold change in luciferase flux from engrafted tumors demonstrated that Pracinostat significantly slowed tumor growth, with additive effects when combined with CD47 blockade (**Figure 5C**). Notably, TAMs isolated from the brains of treated mice exhibited increased phagocytic activity in the Pracinostat cohort, further supporting its role in macrophage reprogramming in vivo. Importantly, the combination with CD47 blockade further enhanced phagocytosis, highlighting the potential synergy between HDAC inhibition and immune checkpoint modulation (**Figure S3B-C**).

## Figures

**Figure S1.**
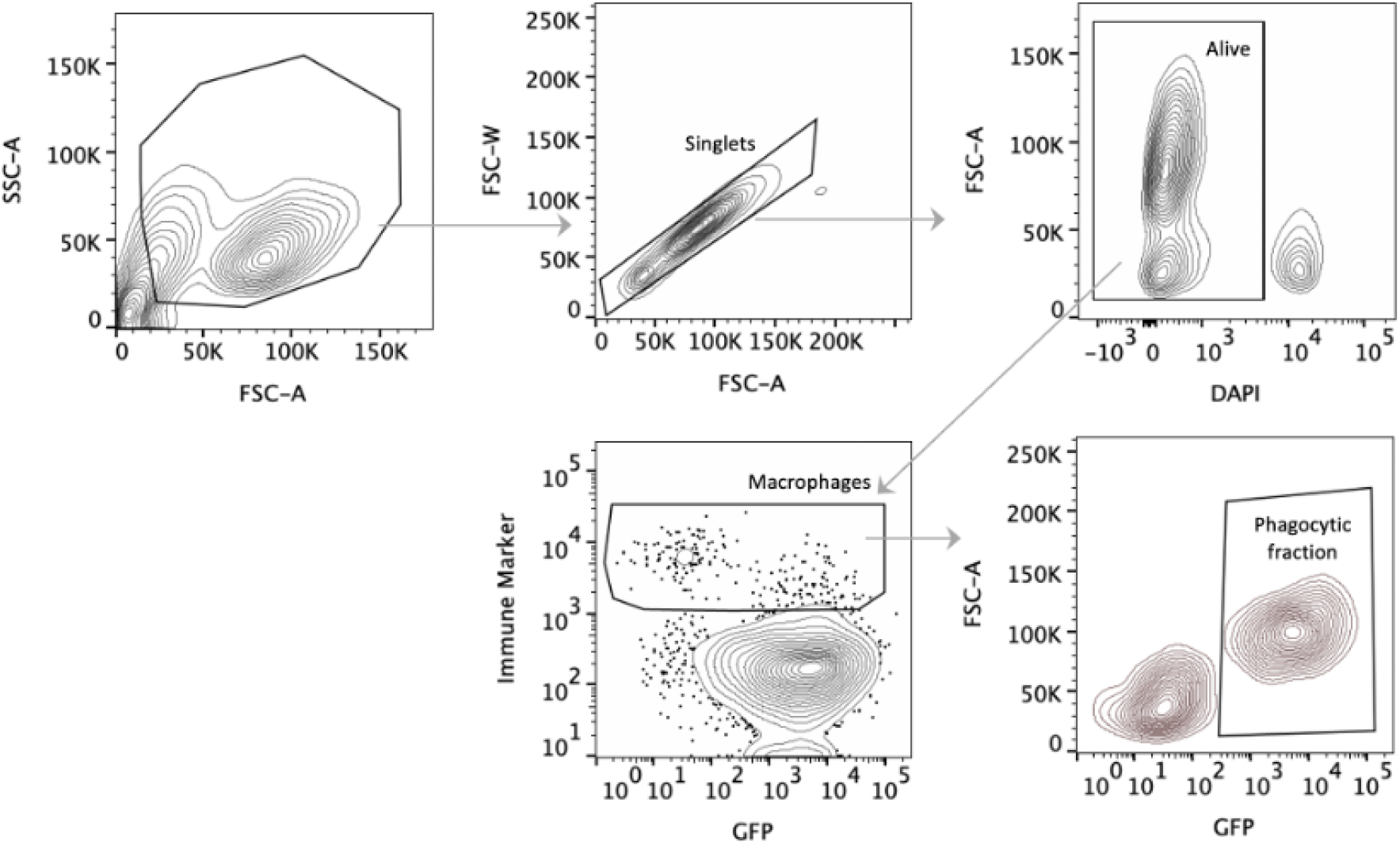
Gating strategy for flow cytometry analysis of primary CD11b+ cells in phagocytosis assays from the chemical screen.

**Figure S2.**
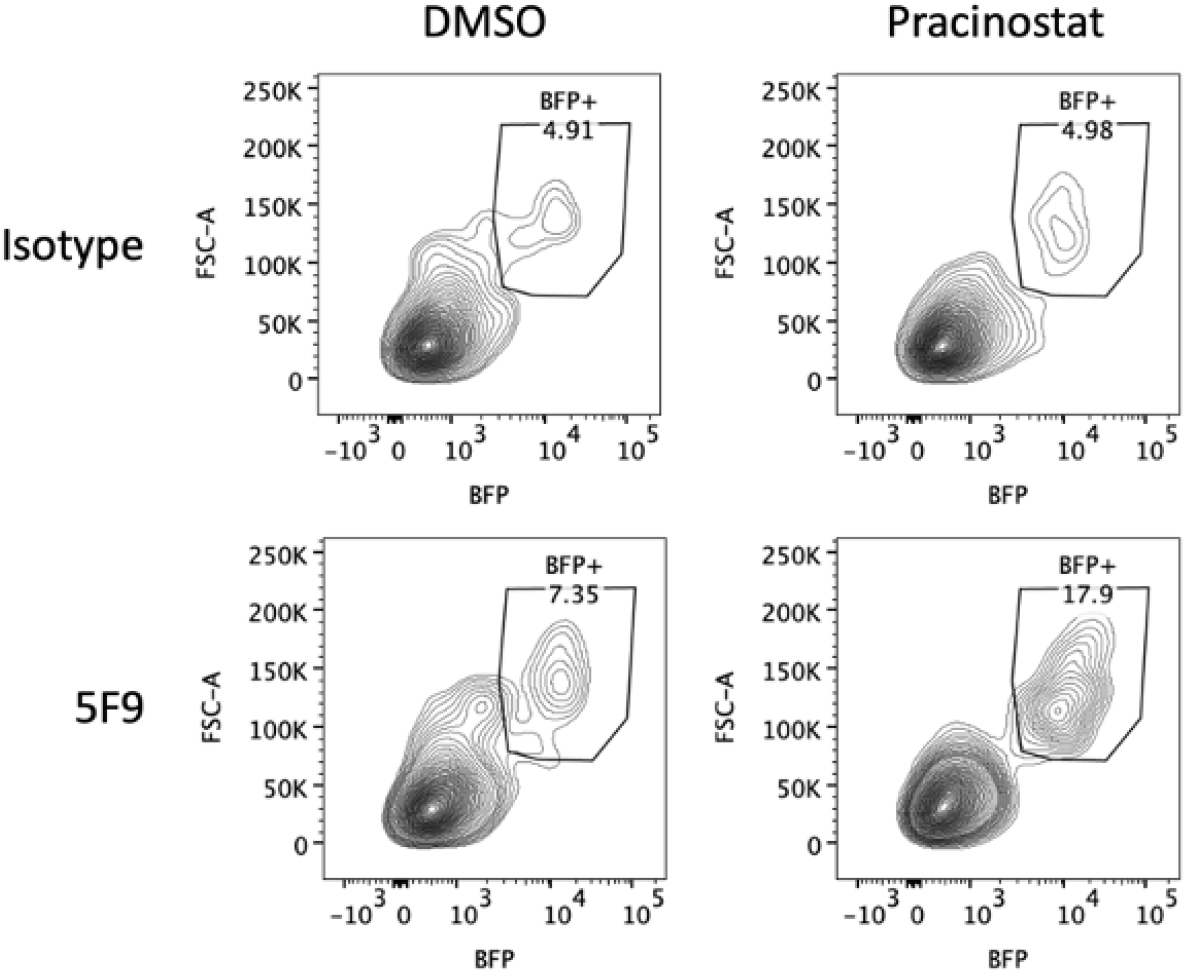
Additional flow cytometry plots of a primary TAM sample from a second patient, demonstrating synergy between Pracinostat and CD47 Abs (Clone 5F9).

**Figure S3.**
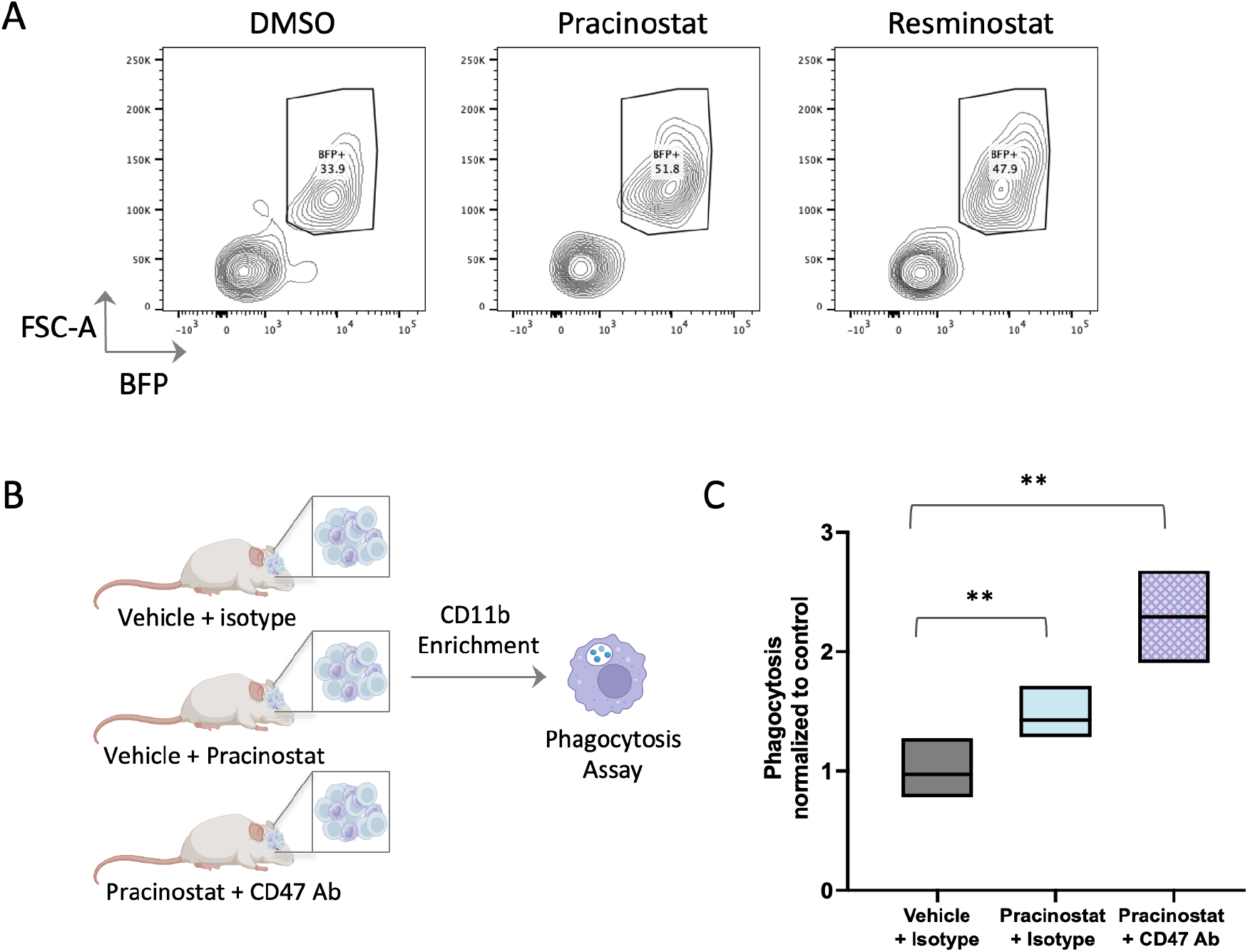
**A**. Flow cytometry analysis of TAMs isolated from a syngeneic GBM mouse model. Phagocytosis was assessed following CD11b+ enrichment and co-incubation with BFP+ T387 GBM cells. Treatment with Pracinostat and Resminostat enhances phagocytosis compared to the DMSO control. **B**. Schematic representation of the workflow for analyzing phagocytosis in a xenograft GBM model. TAMs were isolated from mice four weeks post-surgery, underwent CD11b+ enrichment followed by a two-hour phagocytosis assay. **C**. Quantification of phagocytosis following the assay. Each symbol represents a technical replicate. Pracinostat treatment significantly enhances phagocytosis, with further augmentation observed in combination with CD47 blockade (*, *p* < 0.05). Data are shown as floating boxes, min to max.

## Discussion

This study reports the first high-throughput chemical screen on primary TAMs derived directly from GBM patient samples, and identified FDA-approved drugs that enhance the phagocytic activity of these cells. By using primary microglia and macrophages, rather than immortalized cell lines or *in vitro* differentiated macrophages, we ensured clinical relevance and translatability, addressing a critical gap in GBM drug discovery. Among the compounds identified, HDAC inhibitors, particularly Pracinostat and Resminostat, emerged as potent candidates, consistently enhancing phagocytosis across multiple patient samples. These drugs hold significant promise for repurposing in GBM therapy, especially in combination with CD47-blocking antibodies, which further amplify their effects – consistent with prior findings in melanoma models (24).

While Pracinostat and Resminostat were prioritized for detailed characterization, our screen identified additional hits that remain to be validated. These compounds represent a reservoir of potential therapeutic candidates and warrant further investigation to determine their mechanisms of action and translational potential. This work highlights the potential of high-throughput chemical screening in identifying novel therapeutic strategies for GBM, a disease with few effective treatment options. The inclusion of FDA-approved drugs further accelerates the translational potential of these findings. Future studies will be essential to optimize dosing regimens, assess potential toxicities, and identify patient populations most likely to benefit.

GBM is a highly heterogeneous disease, and TAM composition as well as response to HDAC inhibitors varies between patients. Future studies must evaluate the effect of these agents across different TAM subtypes (e.g., M1/M2 polarization states) and investigate how this could inform precision medicine approaches. Additionally, given the variability in patient responses, identifying predictive biomarkers, such as gene expression profiles, epigenetic markers, or immune signatures, will be crucial for selecting patients most likely to benefit from HDAC inhibitor treatment.

While this study tested the synergy between HDAC inhibitors and CD47 antibodies, there is a strong rationale for exploring other combination strategies to enhance responses to checkpoint inhibitors (e.g., PD-L1 inhibitors) or to complement oncolytic viruses that modulate the tumor microenvironment.

A unique aspect of GBM treatment is that all patients undergo surgical resection, providing a critical window for localized therapeutic interventions. Unlike systemic administration, which may lead to off-target toxicities, delivering HDAC inhibitors and CD47-blocking antibodies locally at the site of tumor excision could enhance therapeutic efficacy while minimizing systemic side effects (25). This localized approach may represent a promising strategy to improve GBM outcomes and warrants further exploration in preclinical and clinical settings.

In conclusion, our study emphasizes the utility of integrating chemical screens to discover and advance new therapies for GBM. By identifying HDAC inhibitors as potent modulators of TAM activity, we have laid the groundwork for a novel approach to GBM treatment. Future efforts will focus on validating additional hits, exploring combination therapies, and advancing these findings toward clinical application to improve outcomes for patients with this devastating disease.

## Materials and Methods

### GBM Cell Culture

The T387 (human patient-derived GBM) and CT2A (murine GBM) cell lines were generously provided by Sam Cheshier’s lab at Stanford University. Cell lines were infected with EF1-EBFP2-T2A-Luciferase lentivirus, double-sorted for EBFP2 signal, and grown in spheres in defined media. Primary GBM cells were generated from a primary GBM sample (IDH wild-type) obtained from a collaboration with Kaiser Permanente. Primary GBM samples used in this study were generously provided by Dr. Gerald Grant (Stanford Neurosurgery) and Dr. Victor Tse (Kaiser Permanente). T387 and primary GBM cells were cultured in maintenance media prepared using a base media comprising Neurobasal media without vitamin A (Invitrogen, 10888022), DMEM/F12 (Invitrogen, 11330-032), nonessential amino acids (Invitrogen, 11140050), sodium pyruvate (Invitrogen, 11360070), 1 M HEPES (Invitrogen, 15630-080), Glutamax (Invitrogen, 35050061), and antibiotic/antimycotic solution (Invitrogen, 15240096). The base media was supplemented with B27 without vitamin A (Life Tech, 12587-010), human EGF (Shenandoah Biotech, 100-26; 20 ng/mL), bFGF (Shenandoah Biotech, 100-146; 20 ng/mL), heparin (Stem Cell Tech, 07980; 2 μg/mL). Cells were seeded in coated T-25 or T-75 flasks and maintained in a humidified incubator at 37°C with 5% CO2, with media changes every 2–3 days. For passaging, cells were detached using Accutase or TrypLE Express, centrifuged at 200 g, and resuspended in fresh media before reseeding. The mouse CT2A cells were cultured in DMEM + GlutaMax (Life Technologies, 10569-010) supplemented with 10% fetal bovine serum (FBS; Life Technologies, 26140079) and 100 U/mL penicillin-streptomycin (Life Technologies, 15140122). Cells were maintained in a humidified incubator at 37°C with 5% CO2 and passaged every 2–3 days at 70–80% confluency. For passaging, cells were detached using TrypLE Express (Life Technologies, 12604013), centrifuged at 300 g for 5 minutes, and reseeded at a 1:3 to 1:5 ratio in fresh media.

### Dissociation of GBM Primary Tumors

Patient-derived tumor-associated macrophages (TAMs) were isolated using a modified tumor dissociation protocol. Tumor tissue was washed in HBSS + Ca/Mg (Thermo Fisher, 14025092) with 1X antibiotic-antimycotic (Thermo Fisher, 15240062) and centrifuged at 200g for 5 minutes at 4°C. The tissue was minced and enzymatically dissociated using collagenase (Sigma-Aldrich, C9407) and DNase (Sigma-Aldrich, DN25) at 37°C with periodic trituration. Residual tissue was further digested with TrypLE Express (Thermo Fisher, 12605010). Red blood cells and debris were removed using ACK lysis buffer (Thermo Fisher, A1049201) and a sucrose density gradient. The suspension was filtered through 100 μm and 40 μm strainers (Corning, 431752 and 431750) and centrifuged at 300g. Final cell suspensions were frozen in BamBanker (VWR, 104085-050). This protocol enabled efficient TAM isolation for downstream applications.

### CD11b enrichment of TAMs

The dissociated tumor samples (mouse or human) were filtered through a 70-µm cell strainer (Corning, 431751) and resuspended in MACS buffer (0.5% BSA and 2 mM EDTA in PBS, Miltenyi Biotec, 130-091-221). CD11b+ cells were isolated using CD11b microbeads (Miltenyi Biotec, 130-097-142) and magnetic separation columns (Miltenyi Biotec, 130-042-201) according to the manufacturer’s protocol. The enriched CD11b+ cells were washed, counted, and immediately used for downstream applications.

### High-throughput drug screening

Patient-derived CD11b+ microglia and macrophages were resuspended in IMDM supplemented with 10% human serum (GemCell, 100-512) at 2.5×10^4^ cells/ml. Cells from the cell suspension were plated, 5×10^3^ cells/well, in ultra-low-attachment 96-well U-bottom plates (Corning, 7007), and treated overnight with 5 µM of individual compounds from a library of FDA-approved chemicals (Selleckchem, L1300), generously provided by the Fattahi lab. The following day, CD11b+ cells were washed in serum-free IMDM media and subjected to phagocytosis assay, described below.

### Phagocytosis Assay

*In vitro* phagocytosis assays were performed by co-culturing GFP-labeled target cells with drug-treated donor-derived macrophages at a ratio of 5-10:1 target cells to macrophages. Target cells were resuspended in serum-free IMDM at 2.5 × 104 cells/well and combined with the overnight-treated CD11b+ cells. The final volume per well was 200 µL. Co-cultures were incubated at 37 °C with 5% CO2 for 1–2 hours. For antibody treatments, 10 µg/mL of antibody (e.g., anti-CD47 or isotype control) was added to target cells before macrophage addition. Co-cultures were incubated for 1–2 hours at 37°C with 5% CO2. Following incubation, cells were stained with anti-CD11b antibody (BioLegend, clone M1/70) or CD45 Ab (Biolegend, clone HI3) for flow cytometry. Phagocytosis was quantified as the percentage of GFP-positive target cells engulfed by macrophages, normalized to controls.

### Orthotopic xenograft models for GBM

a mouse model of GBM was established as previously described (5). Mice were anesthetized in an induction chamber using 3% isoflurane (Minrad International). Once positioned on a stereotactic frame (David Kopf Instruments), anesthesia was maintained at 2% isoflurane delivered via a nose adapter. A burr hole was created 2 mm lateral and 2 mm posterior to the bregma. A blunt-ended needle (Hamilton Co.; 75N, 26s/2-inch point style 2, 5 μL) was inserted through the burr hole to a depth of 3 mm below the dura surface and then retracted by 0.5 mm to create a small reservoir. Using a microinjection pump (UMP-3, World Precision Instruments), 2X105 CT2A or 105 T387-EBFP2-Luc cells were injected in a 3 μL volume at a rate of 30 nL/s. The needle was left in place for 1 minute before being retracted at a rate of 3 mm/min.

### *In vivo* CD47 Ab and Pracinostat Treatment of T387 xenografts

To evaluate the therapeutic efficacy of Pracinostat and CD47 blockade in glioblastoma, we utilized an orthotopic xenograft model. Seven days after implantation of 10^5^ T387 cells, mice were randomized into treatment groups and received one of the following: vehicle + isotype control, vehicle + CD47 Ab (350 µg per dose), Pracinostat (30 mg/kg per dose) + isotype, or Pracinostat + CD47 Ab. CD47 Ab (BioXCell, BE0019-1) was administered intraperitoneally, while Pracinostat was given orally, each four times per week for three weeks. Pracinostat was prepared in a solvent mixture of 10% DMSO (Sigma Aldrich, D2650), 40% PEG300 (MedChem Express, HY-Y0873), 5% Tween-80 (Sigma Aldrich, P1754), and 45% PBS at a final concentration of 3 mg/ml, with each 25 g mouse receiving 250 µl orally (30 mg/kg) via a gavage needle (Fisher, NC1906313). Tumor burden was monitored using bioluminescence imaging (BLI), with tumor progression quantified by measuring the fold change in luciferase flux. At the study endpoint, mice were sacrificed, and tumors were analyzed for macrophage-mediated phagocytosis using flow cytometry.

### TCGA analysis of GBM Patient Survival

Gene expression array data from the TCGA GBM cohort (26) (n=527) were analyzed to assess the prognostic significance of histone deacetylase (HDAC) gene expression. The workflow began by downloading microarray CEL files using the GDC API and extracting normalized gene expression data through Robust Multiarray Average (RMA) normalization using the affy R package. Gene annotations were mapped to the probes using a custom annotation file to generate a final dataset of gene-level expression values. Clinical metadata, including patient survival (days to death), were extracted from XML files available in the TCGA repository and linked to the gene expression data through patient IDs. HDAC7 and HDAC9 were selected as genes of interest, and patients were stratified into high (≥75th percentile) and low (≤25th percentile) expression groups. Kaplan-Meier survival analysis was performed using the lifelines Python package to generate survival curves, with differences between groups assessed using the log-rank test. All analyses were conducted in Python and R, leveraging libraries such as Pandas, Matplotlib, Lifelines, and Scipy for data processing, visualization, and statistical testing. This comprehensive approach integrated large-scale expression data and survival outcomes to provide insights into the role of HDAC expression in GBM prognosis.

### Protein-protein interaction network analysis

Protein-protein interaction network analysis was performed using the Search Tool for the Retrieval of Interacting Genes (STRING) database. Top 40 targets with the highest combined Z scores were used as input for STRING. The minimum required interaction score was set to 0.7, indicating the degree of data support from the following active interaction sources: text mining, experiments, databases, co-expression, neighborhood, gene fusion, and co-occurrence. Cytoscape was used for the visualization of STRING’s interaction output.

## Supporting information

List of compounds tested in the screen, along with their phagocytosis Z-scores and predicted protein targets.

## Acknowledgements

We thank members of the Weissman laboratory:: A.E. Eastman, A.T. Burden, J. Xiang, E. Sedova (Koren) for valuable discussions and brainstorming contributions; T. Naik, A. McCarty for logistical support. We also acknowledge Dr. Gerald Grant (Stanford Neurosurgery) and Dr. Victor Tse (Kaiser Permanente) for providing primary GBM samples, which were essential for this study. The work was generously supported by the NIH/NCI Outstanding Investigator Award (R35CA220434 to I.L.W.); the NIH NIAID (R01AI143889 to I.L.W.); and the Vir the NIH NIAID (R01AI143889 to I.L.W.); the Parker Institute for Cancer Immunotherapy award to I.L.W.; and the Virginia and D.K. Ludwig Fund for Cancer Research (to I.L.W.), as well as NIH Director’s New Innovator Award (DP2NS116769 to F.F) and the National Institute of Diabetes and Digestive and Kidney Diseases (R01DK121169 to F.F.).

